# iNetModels 2.0: an interactive visualization and database of multi-omics data

**DOI:** 10.1101/662502

**Authors:** Muhammad Arif, Cheng Zhang, Xiangyu Li, Cem Güngör, Buğra Çakmak, Metin Arslantürk, Abdellah Tebani, Berkay Özcan, Oğuzhan Subaş, Wenyu Zhou, Brian Piening, Hasan Turkez, Linn Fagerberg, Nathan Price, Leroy Hood, Michael Snyder, Jens Nielsen, Mathias Uhlen, Adil Mardinoglu

**Affiliations:** Science for Life Laboratory, KTH – Royal Institute of Technology, Stockholm, SE-171 21, Sweden; School of Pharmaceutical Sciences & Key Laboratory of Advanced Drug Preparation Technologies, Ministry of Education, Zhengzhou University, Zhengzhou, Henan Province, PR 450001, China; Bash Biotech Inc, 600 West Broadway, Suite 700, San Diego, CA, USA; Department of Metabolic Biochemistry, Rouen University Hospital, 76000 Rouen, France; Normandie Univ, UNIROUEN, CHU Rouen, INSERM U1245, 76000 Rouen, France; Department of Genetics, Stanford University, Stanford, CA 94305, USA; Providence Cancer Center, Oregon Area, Portland; Department of Medical Biology, Faculty of Medicine, Atatürk University, Erzurum, Turkey; Institute of Systems Biology, Seattle, USA; Department of Biology and Biological Engineering, Chalmers University of Technology, Gothenburg, Sweden; Centre for Host–Microbiome Interactions, Dental Institute, King's College London, London, SE1 9RT, United Kingdom

**Keywords:** Multi-omics, network, bioinformatics, systems biology

## Abstract

It is essential to reveal the associations between different omics data for a comprehensive understanding of the altered biological process in human wellness and disease. To date, very few studies have focused on collecting and exhibiting multi-omics associations in a single database. Here, we present iNetModels, an interactive database and visualization platform of Multi-Omics Biological Networks (MOBNs). This platform describes the associations between the clinical chemistry, anthropometric parameters, plasma proteomics and metabolomics as well as metagenomics for oral and gut microbiome obtained from the same individuals. Moreover, iNetModels includes tissue- and cancer-specific Gene Co-expression Networks (GCNs) for exploring the connections between the specific genes. This platform allows the user to interactively explore a single feature's association with other omics data and customize its particular context (e.g. male/female specific). The users can also register their own data for sharing and visualization of the MOBNs and GCNs. Moreover, iNetModels allows users who do not have a bioinformatics background to facilitate human wellness and diseases research. iNetModels can be accessed freely at https://inetmodels.com without any limitation.

## INTRODUCTION

During the past decade, the development of high-throughput technologies has dramatically decreased the cost of generating large-scale multi-omics datasets (1). This has opened up the possibilities to study human wellness and diseases systematically (2). Although analysis of individual omics methodologies has been proven useful in different clinical applications, integrating multi-omics data may offer novel insights and provide a more comprehensive understanding of biological functions in the human body in health and disease (3). For instance, a recent study integrated time series phenomics, metabolomics and fluxomics data from the subjects with various degrees of liver fat, and revealed that non-alcoholic fatty liver disease (NAFLD) is associated with the glycine and serine deficiency (4). Another longitudinal phenomics, transcriptomics, metagenomics, and metabolomics data have been generated for 10 subjects during two weeks follow up study. This study has illustrated the rapid metabolic benefits of an isocaloric carbohydrate-restricted diet on NAFLD patients and revealed the molecular mechanisms associated with the metabolic changes (5). Moreover, several other studies have also demonstrated the benefit of performing longitudinal multi-omics data analysis in systematically capturing human diseases' dynamics (6–8).

To provide a better framework for facilitating these types of investigations, we created iNetModels. This user-friendly platform provides exploratory capabilities and interactive and intuitive visualization of clinical chemistry, anthropometric parameters, plasma proteins, plasma metabolites, oral microbiome and gut microbiome associated with the user-queried features (Figure 1A). The data in iNetModels are obtained from recent studies, where large-scale Multi-Omics Biological Networks (MOBNs) analyses have been performed for individuals with different metabolic conditions. Moreover, we retrieved data from The Cancer Genome Atlas (TCGA) and The Genotype-Tissue Expression (GTEx) Project databases, created normal tissue- and cancer-specific Gene Co-expression Networks (GCNs) and presented the networks in the iNetModels (Figure 1A). The user can simultaneously query for 1-5 features, visualize the selected features and their neighbouring features, download the associated network in both table and figure format and analyse these networks using independent network analysis tools, including Cytoscape (9) and iGraph (10). We also encourage users to upload their networks into iNetModels and make those networks accessible to a broader audience for creating an open platform to share and visualize their networks. To our knowledge, iNetModels is the first database that provides associations between the multi-omics data obtained from the same individuals in a physiological context rather than using a text mining method.

**Figure 1.**
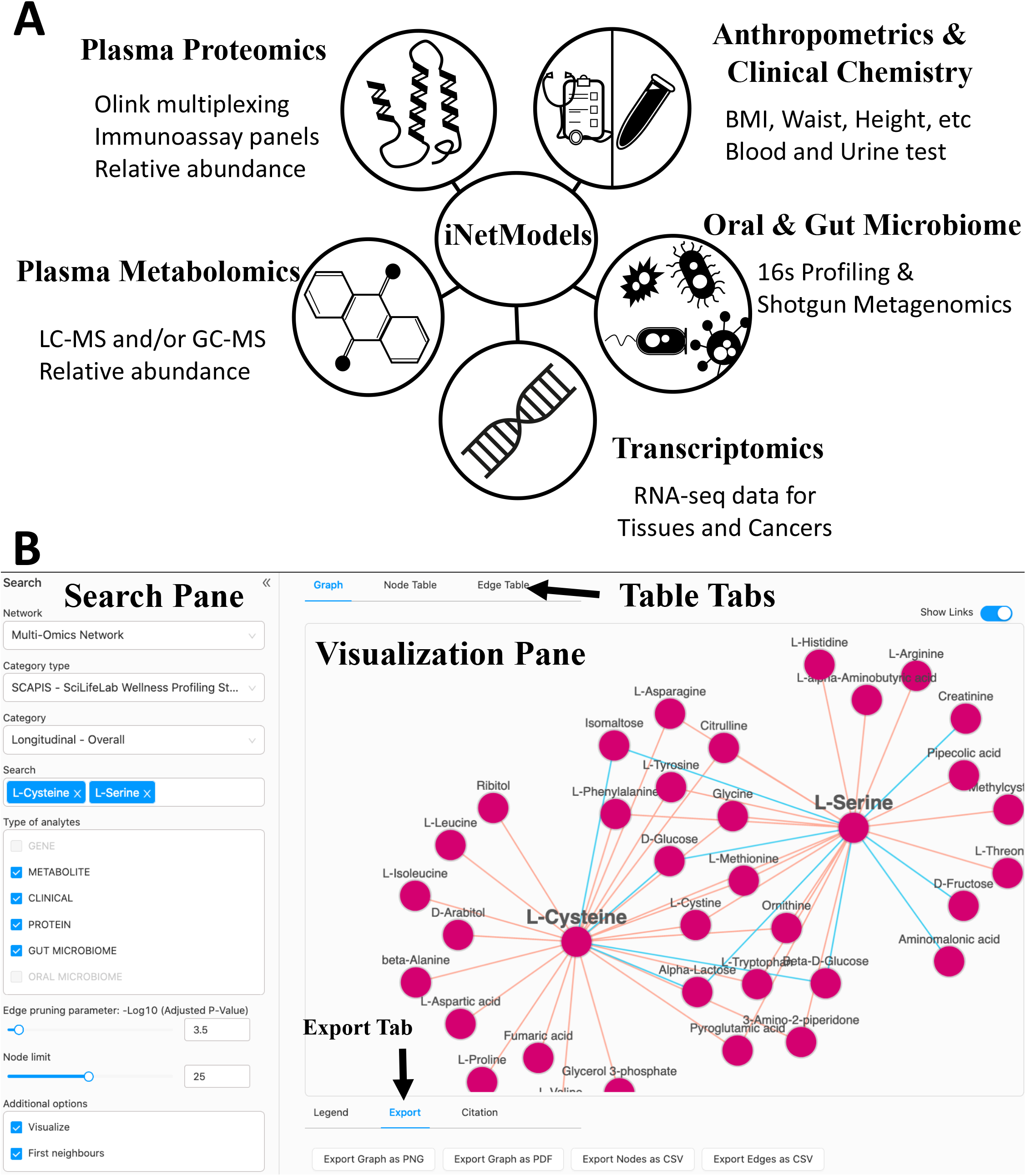
(A) Summary of multi-omics data presented in iNetModels (B) Working page of iNetModels

## PLATFORM DESCRIPTION AND FEATURES

iNetModels 2.0 is a web-based platform that includes two main features: a database and an interactive visualization of multi-omics network analysis (Figure 1B). It is an updated and improved version of the previous work released in 2017 (11). In iNetModels 2.0, we updated the backend and frontend of the platform for a better user experience. We generated more than 100 multi-omics networks, including 85 (tissue- and cancer-specific) GCNs based on TCGA and GTEx databases, and 20 MOBNs, including gender- and insulin-resistant/sensitive-specific (IR/IS) networks, from 3 independent longitudinal wellness profiling studies (12–14) and two different studies, where combined metabolic activators (CMA) are administrated to NAFLD and Covid-19 patients. To our knowledge, this platform is the only publicly available platform that enables exploration of MOBNs that were generated based on personalized data.

### Data Sources and Network Generations

Before generating the MOBNs, we corrected the data by removing the effects of age (in all networks) and sex (in the non-gender-specific networks) using trimmed mean robust regression (13). Moreover, the name of the analytes was standardized in the platform to allow users to compare the relationship between the features across different datasets. MOBNs were built as the consensus of clinical, proteomics, metabolomics, and metagenomics data based on data availability in each study. In SCAPIS-SciLifeLab Wellness Profiling (14) study, clinical, metabolomics, proteomics and gut metagenomics data from 4 visits are presented, whereas in the Integrative Personal Omics (12) and P100 (13) studies, clinical, metabolomics and proteomics data from all visits are presented. For the recently completed CMA supplementation studies, we presented clinical, metabolomics and proteomics data from 2 visits in COVID-19 patients, whereas clinical, metabolomics, proteomics and metagenomics data for oral and gut microbiome from 3 visits in NAFLD patients.

To generate GCNs, we downloaded the data from GTEx and TCGA portals, in transcripts per million (TPM) and fragment per kilobase of transcript per million mapped reads (FPKM) units, respectively. We normalized the TPM and FPKM data and removed the genes with the mean expression below 1 (TPM<1 or FPKM<1) in each tissue or cancer type to avoid bias.

The iNetModels platform provides users a vast number of pre-computed biological networks. Users may choose the suitable networks from tissue- or cancer-specific GCNs to gender or IR/IS-specific MOBNs based on their study's focus. Moreover, in the longitudinal wellness profiling studies, the users can select one of the networks, including cross-sectional and delta networks. These networks are generated based on the methodology presented in the P100 wellness study. Cross-sectional networks represent the multi-omics data correlations in the context of individualized variation. In contrast, delta networks allow users to investigate features that co-vary within the same time intervals.

We combined the data from each omics analysis into a matrix to generate a network and analysed it using the Spearman correlation function from the SciPy package. After filtering for striking correlations between the pairs, we filtered the pairs with FDR < 0.05. Furthermore, we performed community detection analysis using the Leiden algorithm (15) in the iGraph package to identify sub-network clusters for downstream analysis. The number of the nodes and edges in the generated networks varying between 5673 (Liver Cancer) – 12581 (Testis) nodes and 63,292 (Endocervix) – 132,223,286 (Testis) edges in GCNs, whereas 279 (Delta IS) – 1042 (NAFLD CMAs) nodes and 1034 (Delta IS) – 566,390 (SCAPIS – SciLifeLab Longitudinal Male) edges in MOBNs (Supplementary Table 1).

### Features

To search within a specific network (Figure 1B, Supplementary Figures 1), firstly users need to select the specific network category (GCNs or MOBNs), then select the specific network type (normal tissue, cancer, or multi-omics study), and subsequently select the specific network. Following that, users need to input the commonly known names of analytes (gene, protein, metabolite name etc.) of interests using the free text and/or drop-list. Optionally, users can filter the network based on the analyte types (in multi-omics networks) and statistical properties, e.g. FDR and Spearman correlation ranking (absolute values, limited to top 50 with visualization). Furthermore, users can filter the networks based on first neighbours' connectivity if they just want to visualize or download the complete tables without visualization.

Once the network is generated, the users can interactively explore the network, download the network as a figure and its content in a table format for further exploration and downstream analysis. All information about each analyte (network nodes) and related associations (network edges) is shown in the table area: the analyte name, short description, unit, correlation and the P-value of the significant associations etc. Besides, wherever possible, analytes are linked with external databases such as KEGG (16), Human Protein Atlas (17–19), Uniprot (20), and HMDB (21) to facilitate further biological interpretation and investigation. All of this information can be downloaded directly and is compatible with other network analysis tools and software, such as Cytoscape or iGraph package in Python and R. By using these tools, users can merge multiple networks and perform additional downstream analysis.

Moreover, in the iNetModels 2.0, we implemented programmatic access to the database using an in-house-built Python package that can be found under the “API” section. We currently limit the query to 1 query per second.

### Case Study related to administration of CMA in NAFLD

One of our platform's unique feature and superior strength is that iNetModels 2.0 is the first and only platform that supports the exploration of MOBNs based on personalized data. This is a great advantage to avoid bias since all data were analysed in a paired manner.

In our recent study (22), we tested a potential therapeutic strategy for NAFLD patients through CMA administration. We provided CMA to 10 subjects involved in the trial and collected plasma samples during the day to generated proteomics and metabolomics data. The data generated in the clinical trial were analysed using metabolic modelling. The results of the analysis were validated by performing animal experiments, where l-serine was supplemented to mouse and a reduction in the liver triglycerides (TG) and markers of liver tissue functions, e.g. ALAT, ASAT, and ALP was observed. We validated these results in two independent MOBNs in iNetModels 2.0 (Figure 2A, Supplementary Figures 2A). Our analysis revealed that two metabolic activators including L-Serine and L-Cysteine are positively associated with metabolites related to branched amino-acid metabolism (i.e. L-Valine, L-Isoleucine, and L-Leucine) and negatively associated with plasma glucose level, supporting the main findings of the study (Figure 2B). The MOBNs also showed that l-serine is associated negatively with cholesterol-related clinical variables (ApoB, TG) and several other inflammation markers (hsCRP, IL1RN, and CD300C).

**Figure 2.**
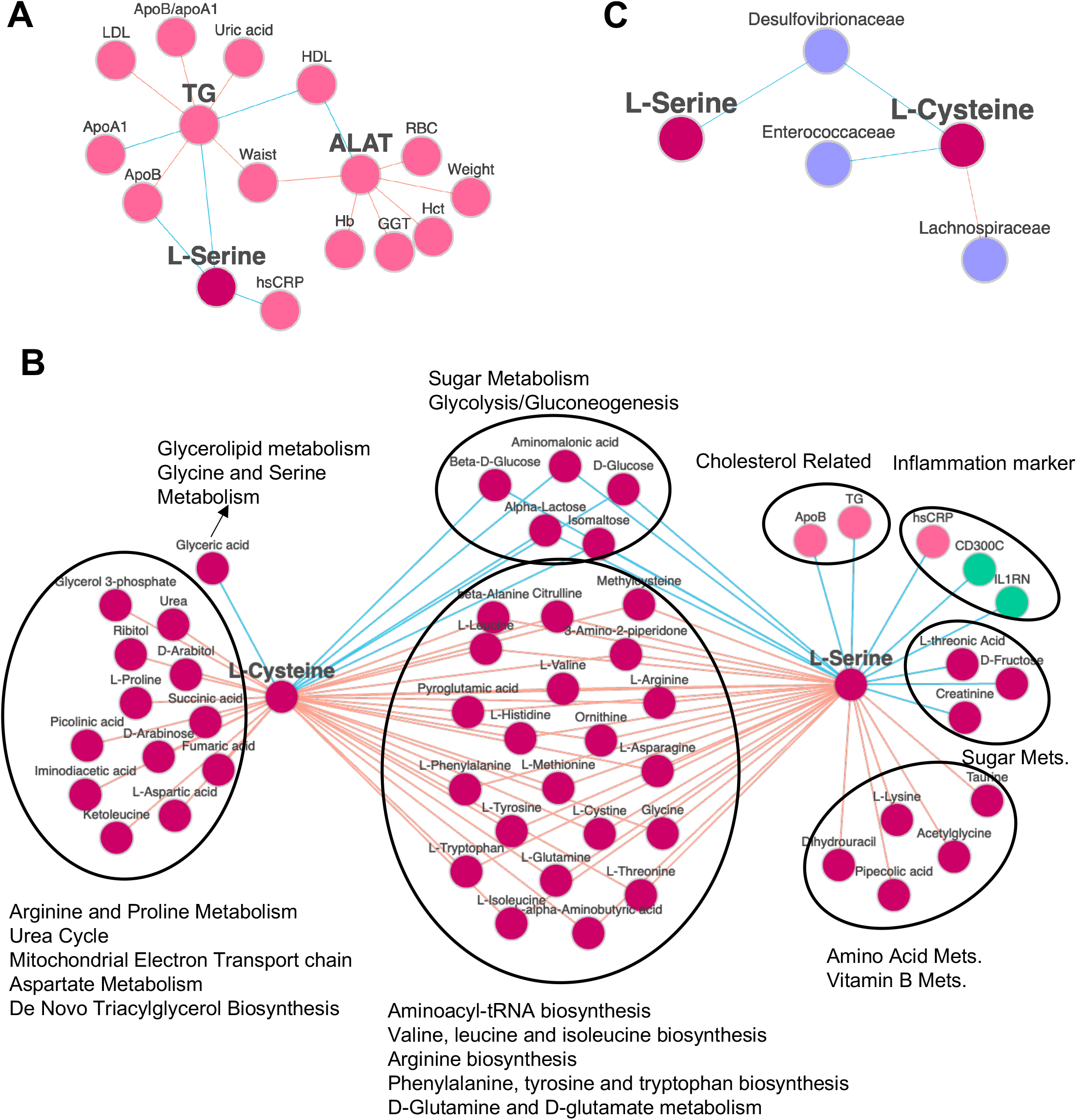
(A) Validation of the hypothesis about the supplementation of L-Serine that was associated with the decrease in the plasma triglycerides levels and liver enzyme (ALAT) in the SCAPIS-SciLifeLab Wellness Profiling study. (B) The two main components of the supplementation (L-Cysteine and L-Serine) and their neighbouring analytes are presented based on multi-omics biological networks analysis (C) The two main components of the supplementation (L-Cysteine and L-Serine) and their associations with the species in the gut microbiome are presented. All networks in this figure were taken from cross-sectional overall SCAPIS-SciLifeLab Wellness Profiling study.

Based on the same network, we can filter to show only specific analyte types (Figure 2C, Supplementary Figures 2B-D). For example, we used the same network to show the association of L-Cysteine and L-Serine with only gut microbiome (Figure 2C), as dysbiosis in the gut microbiome has been associated with NAFLD (23,24). We observed that L-Cysteine had a positive correlation with the abundance of Lachnospiraceae and negative correlation with the abundance of Enterococcaceae family. Independent studies have shown that the abundance of Lachnospiraceae was increased in cirrhosis (25,26), whereas the abundance of Enterococcaceae was decreased in cirrhosis (25). We also found that the levels of both L-Serine and L-Cysteine were negatively correlated with the abundance of Desulfovibrionaceae family, which was increased with the severity of NAFLD (27).

## CONCLUSION

iNetModels is a unique platform that gathers a broad spectrum of biological networks, from tissue- and cancer-specific GCNs to MOBNs based on personalized data. This platform may help researchers to perform exploration and validation experiments, identify functional relationships between the analytes, and most importantly, provide new insights into the biological experiments and ultimately identify potential drug targets and biomarkers. The case study about the supplementation of CMA in NAFLD patients has shown the efficient usage of this platform in testing and validating hypotheses as well as confirming results from experiments or clinical trials. This platform is designed in a user-friendly way and it is freely accessible to a wide range of users including bench scientists with limited or no formal bioinformatics background. We also envisage that this platform will be a key resource for computational biologists working in omics data integration, network science, systems biology and systems medicine. In this context, we expect iNetModels to be an essential resource for more in-depth multi-omics analysis that may reveal novel molecular mechanisms underlying human wellness and disease.

## AVAILABILITY

iNetModels can be accessed freely by everyone at https://inetmodels.com without any limitation. All codes used to generate the network and the network data are available under the “API” and “Help” section of the website.

## SUPPLEMENTARY DATA

Supplementary Data are available at NAR Online.

## ACKNOWLEDGEMENT

We appreciate the data sharing from dbGaP. This work was supported in part by the Robert Wood Johnson Foundation, the M.J. Murdock Charitable Trust, NIH grants 2P50GM076547, ES017885, RC2HG005805, and Arivale. The Genotype-Tissue Expression (GTEx) Project was supported by the Common Fund of the Office of the Director of the National Institutes of Health, and by NCI, NHGRI, NHLBI, NIDA, NIMH, and NINDS. The data used for the analyses described in this manuscript were obtained from the GTEx Portal on 10/21/20. The results shown here are in part based upon data generated by the TCGA Research Network: https://www.cancer.gov/tcga. The computations were performed on resources provided by SNIC through Uppsala Multidisciplinary Center for Advanced Computational Science (UPPMAX) under Project SNIC 2019/8-259.

## FUNDING

This work has been supported by the Knut and Alice Wallenberg Foundation and Bash Biotech Inc, San Diego, CA, USA.

## CONFLICT OF INTEREST

Cem Güngör, Buğra Çakmak, Metin Arslantürk, Berkay Özcan, Oğuzhan Subaş are the employees of Bash Biotech Inc. The other authors declare no conflict of interest.

---

**Supplementary Figures 1.**
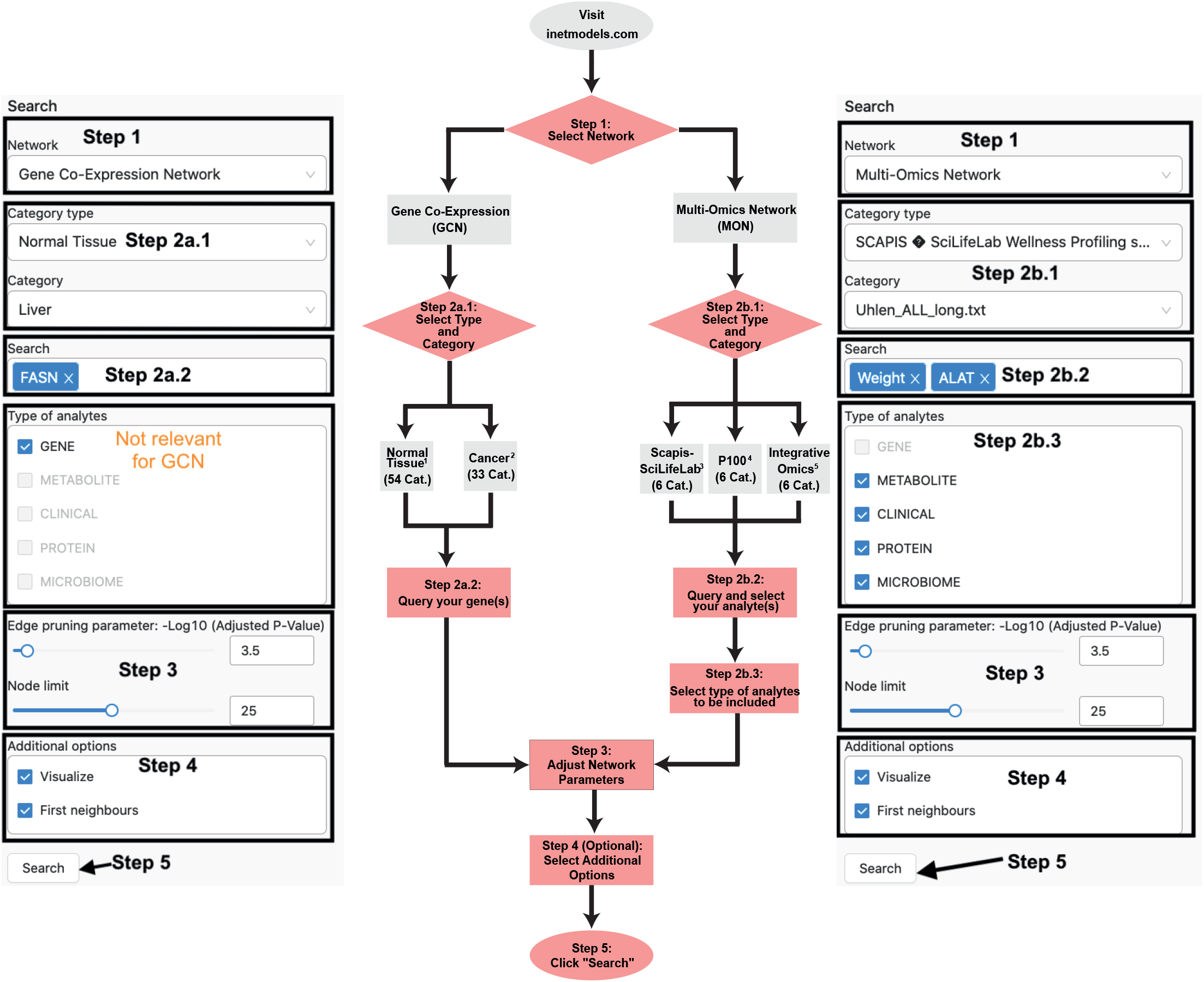
User query flow in iNetModels to search for specific analytes of interest.

**Supplementary Figures 2.**
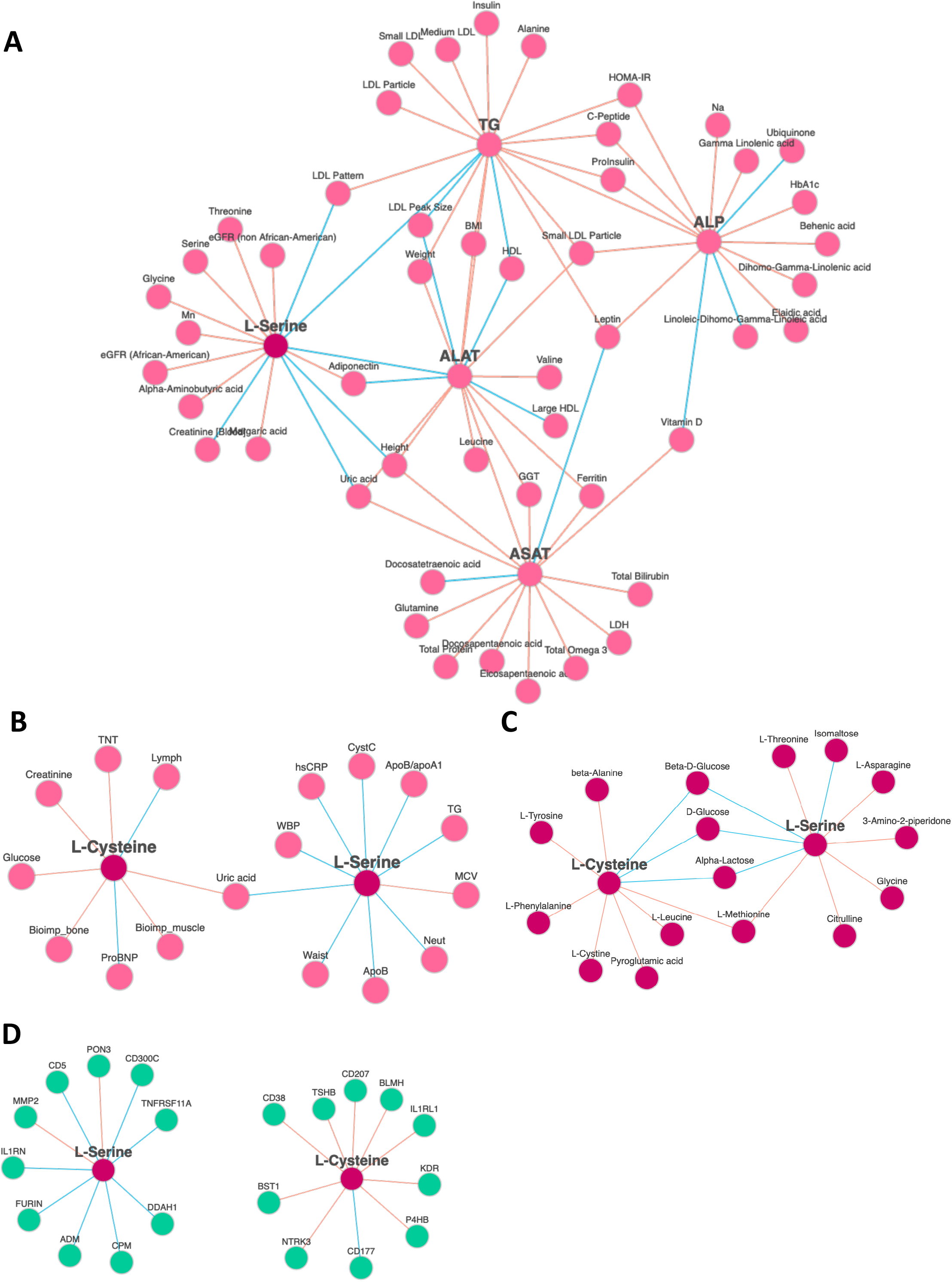
(A) Validation of the hypothesis about the supplementation of L-Serine that was associated with the decrease in the plasma triglyceride levels and liver enzymes (ALAT, ASAT, and ALP) in the P100 Study. (B) Clinical data, (C) Metabolites, and (D) Proteins associated with the two main components of the supplementation (L-Cysteine and L-Serine) based on multi-omics biological networks analysis in the SCAPIS-SciLifeLab Wellness Profiling Study.

**Supplementary Table 1** Detailed individual network information, including the number of node and edges.

## Notes

https://inetmodels.com

### Competing Interest Statement

Cem Gungor, Bugra Cakmak, Metin Arslanturk, Berkay Ozcan, Oguzhan Subas are the employees of Bash Biotech Inc. The other authors declare no conflict of interest.

### Summary of Updates

Expanding the paper; adjusting the format to comply with NAR webserver issue; adding a use case; adjustment of authorship + their affiliations; adding supplemental files; figures updated

